# IL-22BP production is heterogeneously distributed in Crohn’s disease

**DOI:** 10.1101/2021.04.07.438837

**Authors:** Aurélie Fantou, Eric Lagrue, Thomas Laurent, Laurence Delbos, Stéphanie Blandin, Anne Jarry, Gaëlle Beriou, Cécile Braudeau, Nina Salabert, Eros Marin, Aurélie Moreau, Juliette Podevin, Arnaud Bourreille, Régis Josien, Jérôme C Martin

## Abstract

Crohn’s disease (CD), a form of inflammatory bowel disease (IBD), is characterized by impaired epithelial barrier functions and dysregulated mucosal immune responses. IL-22 binding protein (IL-22BP) is a soluble inhibitor regulating IL-22 bioactivity, a cytokine proposed to play protective roles during CD. We and others have shown that IL-22BP is produced in IBD inflamed tissues, hence suggesting a role in CD. In this work, we extended the characterization of IL-22BP production and distribution in CD tissues by applying enzyme-linked immunosorbent assays to supernatants obtained from the culture of endoscopic biopsies of patients, and reverse transcription-quantitative polymerase chain reaction on sorted immune cell subsets. We reveal that IL-22BP levels are higher in inflamed ileums than colons. We observe that in a cell-intrinsic fashion, populations of mononuclear phagocytes and eosinophils express IL-22BP at the highest levels in comparison to other sources of T cells. We suggest the enrichment of intestinal eosinophils could explain higher IL-22BP levels in the ileum. In inflamed colon, we reveal the presence of increased IL-22/IL22BP ratios compared to controls, and a strong correlation between IL-22BP and CCL24. We identify monocyte-derived dendritic cells (moDC) as a cellular subtype co-expressing both cytokines and validate our finding using *in vitro* culture systems. We also show that retinoic acid induces the secretion of both IL-22BP and CCL24 by moDC. Finally, we report on higher IL-22BP levels in active smokers. In conclusion, our work provides new information relevant to therapeutic strategies modulating IL-22 bioactivity in CD, especially in the context of disease location.

## Introduction

Inflammatory bowel disease (IBD) is a group a chronic inflammatory conditions of the gastrointestinal tract, the main clinical entities of which are Crohn’s disease (CD) and ulcerative colitis (UC) (1). IBD results from dysregulated mucosal immune responses against the microbiota, triggered by environmental factors in genetically susceptible individuals (2). Inflammatory responses in IBD alter the intestinal epithelial barrier, further amplifying immune pathogenicity. The cytokine interleukin-(IL) 22 is proposed to play central roles during IBD inflammation (3,4). IL-22 is induced during IBD flares, as a result of increased production by CD4^+^ T cells and group 3 innate lymphoid cells (ILC3) (5). IL-22 receptor (IL-22R) expression is mostly limited to epithelial cells and protective functions for IL-22 on the gut epithelium have been suggested in several rodent models of intestinal inflammation (6,7). IL-22 supports gut epithelial barrier properties by inducing the secretion of antimicrobial peptides (6) and mucins (7), as well as intestinal epithelial cell (IEC) survival and proliferation (8). When dysregulated, however, IL-22 actions on IEC can promote tumor cell proliferation (9,10). IL-22 binding protein (IL-22BP ; encoded by the gene *IL22RA2*) is a soluble inhibitor specific for IL-22 (11), which regulates the level of IL-22 bioactivity *in vivo* (12,13). Concordantly, we have observed increased IL-22-dependent protection against DSS-induced colitis in IL-22BP-deficient rats (14). In mice, IL-22BP worsens T cell-dependent colitis (15) but prevents tumorigenic long-lasting pro-proliferative actions of IL-22 (12). We and others have shown that the expression of *IL22RA2* is down-regulated during infectious colitis but not in IBD inflamed tissues, hence suggesting possible pathophysiological relevance for the IL-22BP-dependent modulation of IL-22 bioactivity (14,15). IL-22BP is produced by various cell types, which include subsets of intestinal dendritic cells (DCs) and macrophages in the gut lamina propria (LP) and secondary lymphoid structures (12,15–18). In human, we have revealed that IL-22BP is also produced by gut eosinophils in the LP of both healthy and inflamed IBD tissues (14). Finally, CD4^+^ T cells have also been proposed to contribute to IL-22BP levels detected in IBD (15,19).

Several layers of disease heterogeneity exist in CD. It has so far remained unclear whether the general findings about IL-22BP regulation in IBD discussed above can be extrapolated homogenously to patients with CD, especially when considering intestinal segments of disease location. In the light of therapeutic strategies modulating IL-22 bioactivity in IBD that are currently under evaluation, we thus sought to extend further the characterization of IL-22BP production and distribution in intestinal tissues from a cohort of CD patients.

## Material and Methods

### Patients

Patients were recruited, from January 2016 to October 2019, from the Institut des Maladies de l’Appareil Digestif (IMAD), the gastroenterology department and the digestive endoscopy unit at CHU Nantes Hospital. The protocol was approved, and informed consent was obtained from all participants in accordance with the institutional review board (DC-2008-402). A cohort 1 of CD patients was constituted to explore cytokine secretions in supernatants from *ex vivo* biopsy cultures. Cohort 1 included a total of 43 CD patients for whom endoscopic biopsies in involved areas were collected during colonoscopies planned for routine care (**Supplementary Table 1**). Control biopsies were also analyzed and collected from surgical specimens of colon cancers, on unaffected areas of the colon or the ileum, at least 10 cm distant from the tumor. A cohort 2 that included mesenteric lymph nodes and intestinal mucosa obtained from ileocecal resections of 6 patients with active CD was constituted for gene expression analyses in sorted immune cell subsets (**Supplementary Table 2**).

### *Ex vivo* cultures of gut biopsies

During the colonoscopy procedure, biopsies were collected with forceps directly in ice cold RPMI and processed within 2h. Biopsies (controlled for weight) were put into 4-well Petri dishes filled with in 500μL serum-free medium (RPMI 1640, Gibco™) supplemented with BSA (0.01%), 200μg/mL Penicillin/Streptomycin (Gibco ™; ref 15140-122) and 0.25μg/mL Fungizone (Gibco™; ref 15290-026), and cultured *ex vivo* during 6 h at +37°C in a 95% O_2_ / 5% CO_2_ atmosphere on a low-speed rocking platform. Supernatants were collected and stored at -80°C until use.

### ELISA assay for 22BP

IL-22BP quantification was performed with the Human IL-22BP ELISA DuoSet kit (R&D System, ref DY-1087-05) according to manufacturer’s instruction. Briefly, a 96-well microplate was coated with a rabbit monoclonal anti-IL-22BP [4μg/mL] and incubated at room temperature (RT) overnight. The next day, after blocking with Reagent Diluent (R&D System, ref DY006), 100-μL standard dilutions and samples were added to each well and incubated 2 hours at RT. Then, 100μL/well of goat anti-IL-22BP [70ng/mL] were added and 2 h incubation at RT was performed. Finally, ELISA was revealed and plates were read at 450 nm with TECAN Spark® instrument.

### Multiplex assay

Levels of soluble cytokines (CCL4, CXCL10, CXCL2, CCL24, CCL11, IFNγ, IL-10, IL-17A, IL-22, IL-27, IL-6, IL-8, CCL2, OSM, CXCL1, CXCL11, CXCL5, CCL26, IL-1β, IL-18, IL-23, IL-33, IL-7, CXCL9, TNF) were quantified in supernatants obtained after *ex vivo* biopsy cultures with a multiplex assay from Biotechne (Rennes, France) and a Luminex MAGPIX® instrument (**Supplementary Table 3**).

### Eosinophil Counts

Eosinophil counts were scored in a blinded fashion by a trained pathologist. Hematoxylin and eosin-stained slides from intestinal tissues of 36 CD patients were analyzed with an Olympus BH2 microscope. A high-power field (HPF) included an area of 0.196mm^2^. For each slide, eosinophil counts were defined by averaging the results obtained in 5 different HPFs selected randomly. Results of eosinophil counts were reported as the number of eosinophils/mm^2^.

### Isolation of intestinal lamina propria cells

Tissues from surgical resections were collected in ice cold RPMI 1640 (Gibco™) and processed within one hour after the end of the surgery. The mucosa was stripped and cut into small pieces before transfer into complete RPMI media. Epithelial cells were dissociated in an EDTA-enriched dissociation medium (HBSS w/o Ca^2+^ Mg^2+^ (Gibco™) - HEPES 10mM (Gibco™) - EDTA 5mM at +37°C through two 15 min cycles of agitation at 100 rpm. After each cycle, intestinal fragments were hand-shaken for 30 s and vortexed vigorously for another 30 s. Epithelium-free fragments were washed in PBS, and transferred in the digestion medium (HBSS with Ca^2+^ Mg^2+^ - Fetal Calf serum (FCS) 2% - DNase I 0.1mg/mL (Sigma-Aldrich, ref 11284932001) – Collagenase IV 0.5mg/mL (Sigma-Aldrich, ref C5138) for 40 min at +37°C under 100 rpm agitation; and vortexing every 20 min. Cell suspensions were filtered through 70μm pore size cell strainers (BD Biosciences) and washed with FACS-buffer (with EDTA 1mM) twice before being processed for cell sorting.

### Intestinal mesenteric lymph nodes isolation

Tissues were collected in ice cold RPMI 1640 (Gibco™) from the resected specimens and processed within one hour after the end of the surgery. After removing the adipose tissue, lymph mesenteric nodes were cut in small pieces and transferred into the digestion medium (10mL of complete RPMI with Collagenase D (2mg/mL-Sigma-Aldrich, ref 1108882001) and DNase I (0,1 mg/mL)). After 30 min of incubation at +37°C 100 rpm, EDTA (1mM) was added to block the reaction. Cell suspensions were filtered through 70μm pore size cell strainers (BD Biosciences) and processed for cell sorting.

### Cells sorting

Single cell suspensions were incubated in PBS containing the flow cytometry antibody cocktail (**Supplementary Table 4**) for 20 min at 4°C in the dark. Dead cells were excluded by gating on 4’,6-diamidino-2-phenylindole (DAPI)-negative cells. Cell sorting was performed on a BD FACS Aria Cell sorter (BD Biosciences) using the gating strategies shown in Figure 2.

### Real-time quantitative PCR

Sorted cells were suspended in TRIzol reagent (Thermo Fisher Scientific) and frozen at -80°C. Total RNA was extracted using an RNeasy mini kit (Qiagen,Valencia, CA) according to manufacturer’s instructions. Reverse transcription was performed using Murine Moloney Leukemia Virus Reverse Transcriptase (Thermo Fisher Scientific) following manufacturer’s instructions. Gene expressions were assessed with the TaqMan Fast Advanced Master Mix reagent (Applied Biosystems, Foster City, Calif). Primers and probes were from Applied Biosystems (**Supplementary Table 5**). Real-time PCR was performed using the StepOne Plus System (Applied Biosystems). Relative expression was normalized to hypoxanthine-guanine phosphoribosyltransferase and calculated using 2^^-ddct^ method. Results were expressed in arbitrary units (a.u.).

### Monocyte-derived dendritic cells

Monocytes from healthy volunteers were isolated either by elutriation of PBMCs (Clinical Development and Transfer Platform, Nantes, France) or by magnetic labelling (untouched cells, Human monocyte Isolation kit II). To obtain monocyte-derived DC (moDC), 2.5×10^6^ monocytes were incubated in a 6-well plate in 5 mL of complete medium (RPMI 1640 medium containing 10% FCS, 1% L-glutamine, 1% antibiotics, 1mM Sodium Pyruvate, 1mM HEPES, 1% non-essential amino acids) supplemented with recombinant human IL-4 (200U/mL) and recombinant human GM-CSF (100U/mL) for 6 days at 37°C with 5% CO_2_. When indicated, cells were treated with retinoic acid (RA) (100nM, Sigma Aldrich), LPS (1 μg/mL, Sigma Aldrich) and TNF (50 ng/mL, Miltenyi). After 6 days, moDC and supernatants were collected and frozen at −80°C until use.

### Immunofluorescence staining from mesenteric lymph nodes

Lymph nodes isolated from 3 CD patients were frozen in tissue-Tek. Sections were fixed for 15 min in paraformaldehyde. After rehydration, a 10 min step saturation was performed with H2O2 (3%). For double staining, mouse IgG anti-hIL-22BP (MAB 1087 from R&D System, [1/800]) or isotype control (mouse IgG1, DDXCMO1P from Dendritic products [1/800] were incubated at room temperature (RT) for 1 hour. After washing, several steps are carried out in order to amplify the IL-22BP signal: polymer enhancer (M023 from ImPath) then HRP-2 (M024 from ImPath) and finally Opal 650 (FP1496A from Akoya) following manufacturer’s instruction. Then, after washing and a second step of saturation with goat serum 1.5%, a second purified antibody: rabbit anti-hCD3 (A0452 from Dako, [1/800]) or isotype control rabbit IgG (I-1000 from Vector) was added and incubated at RT during 1 hour. Purified antibody was revealed with adapted secondary antibody labelled with Alexa488 (goat anti-rabbit, A11008) from Life Technologies. After washing, DAPI (Molecular Probes, D1306) was incubated 30 min. Slides were mounted with Vectashield® Vibrance™ Antifade Mounting Medium (H-1700 Vector Laboratories). Images were obtained with A1 R Si Confocal microscope (MicroPICell).

### Cytokine correlation plots

Pairwise correlations between cytokines were calculated and visualized as a correlogram using R function corrplot. Spearman’s rank correlation coefficient (ρ) was indicated by heat scale; significance was indicated by *P < 0.05, **P < 0.01, and ***P < 0.001.

### Statistical analysis

Statistical analysis was performed with GraphPad Prism Software (GraphPad Software, San Diego, CA). Means comparisons of unpaired samples were performed using the Mann–Whitney U-test or the Kruskal–Wallis test with Dunn’s post-test. The Wilcoxon signed-rank test was used for paired samples. P-values <0.05 were considered statistically significant.

## Results

### IL22BP levels are higher in the ileum than in the colon of Crohn’s disease patients

Both IL-22 and IL-22BP mediate their biological functions as secreted soluble proteins (11,20). To explore IL-22BP production in a biologically relevant manner in CD, we thus quantified secreted levels released after short *ex vivo* culture of intestinal biopsies. Soluble IL-22BP was consistently detected in supernatants from both controls and CD biopsies (see **Supplementary Table 1** for patient characteristics), and though means were not statistically significant, the highest levels were detected in a subset of CD patients (**Figure 1A**). To begin explore what factors would preferentially associate with IL-22BP heterogenous distributions in CD, we ran a multiple linear regression analysis that included several disease-relevant clinical parameters such as age, sex, disease location, disease phenotype and medication (**Table 1**). This analysis revealed disease location as the only factor reaching statistical significance (Student t-test t=2.693; P=0.012). Accordingly, higher levels of IL-22BP were detected in the supernatants of ileal vs. colonic CD biopsies (**Figure 1B**). IL-22 levels in turn, were similar between biopsy supernatants from both locations, which translated into IL-22/IL-22BP ratios, a surrogate to infer IL-22 bioactivity, higher in the colon as compared to the ileum of CD patients (**Figures 1C&1D**). In fact, the analysis of IL-22BP and IL-22 in control patients indicated that both proteins were produced at higher levels in ileal vs. colonic biopsies (**Figures 1E&F**), an observation confirming previous reports about the physiological distribution of IL-22 across intestinal segments (21). Remarkably, however, IL-22 induction was largely limited to CD colons (**Figures 1G&H**). IL-22BP mean levels in turn, remained similar between controls and CD patients when controlling for gut segments (**Figures 1I&J**). Concordantly, in comparison to controls, IL-22/IL-22BP ratios were unchanged in the ileum but increased significantly in CD colons (**Figures 1K&L**).

**Table 1:** Multiple linear regression to assess associations between clinical parameters of CD and IL-22BP levels.

| Variable | \|t\| | P value |
| --- | --- | --- |
| Sex | 0.7929 | 0.43 |
| Age | 0.0998 | 0.92 |
| Localization | 2.693 | **0.012** |
| Tobacco | 1.813 | 0.08 |
| Disease evolution | 0.0809 | 0.94 |
| Phenotype | 0.0845 | 0.93 |
| Perianal involvement | 0.8205 | 0.42 |
| Treatment: anti-TNF | 0.4463 | 0.66 |
| Treatment: IS | 0.5842 | 0.56 |
| Surgical resection | 1.108 | 0.28 |
| Clinical score | 0.033 | 0.97 |

**Figure 1:**
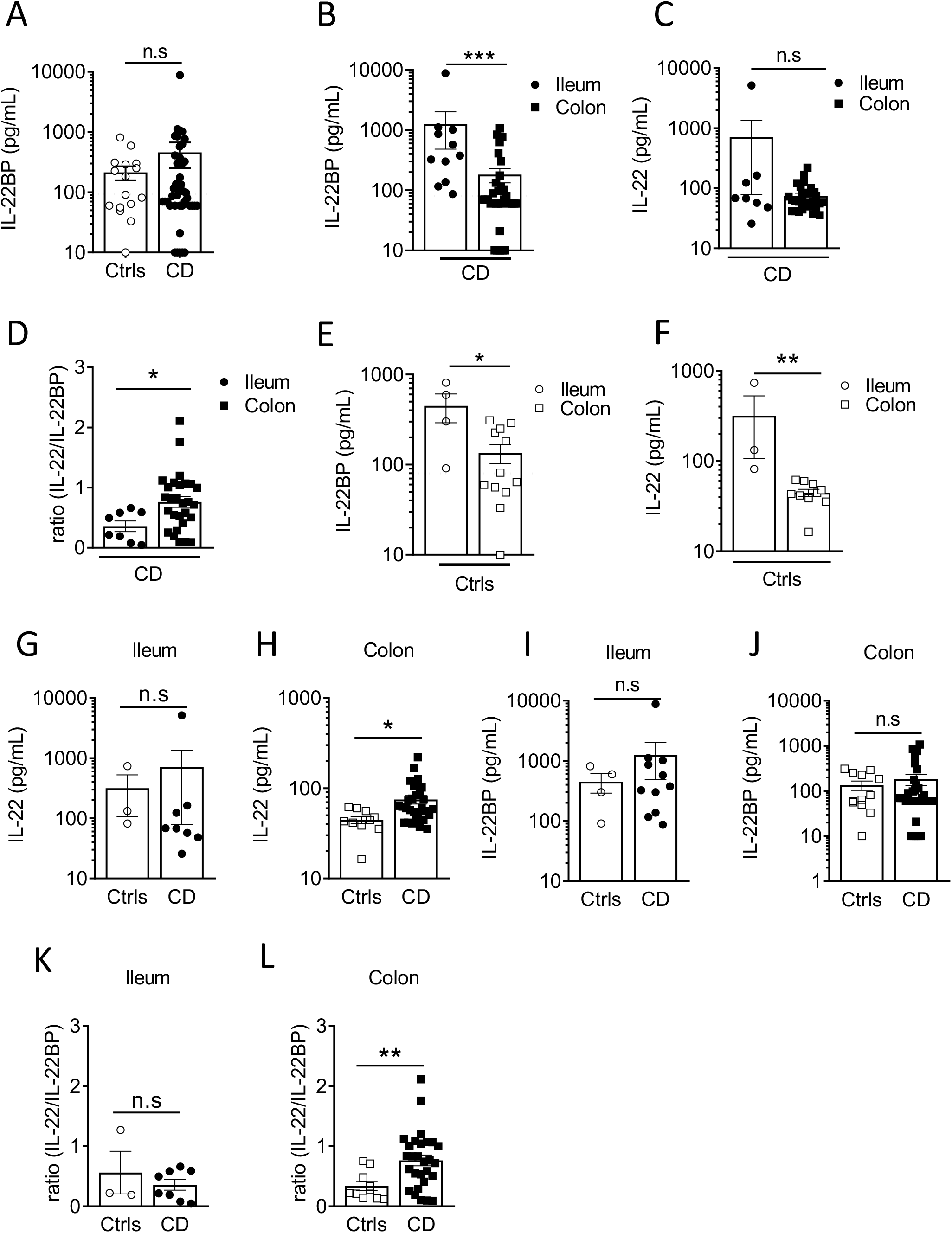
IL22BP levels are higher in the ileum than in the colon of Crohn’s disease patients. (**A**) Levels of soluble IL-22BP were quantified by ELISA in the supernatant of gut endoscopic biopsies from controls (Ctrls) (n=16) and Crohn’s disease (CD) (n=43) patients after 6 hours of ex vivo culture. (**B-C**) Quantification of soluble IL-22BP (B) and IL-22 (C) in culture supernatants of biopsies from CD patients with inflamed lesions in the ileum (n=11) or in the colon (n=32). (**D**) Comparison of IL-22/IL-22BP ratios in the supernatants of CD patients with inflamed ileum and colon. (**E-F**) Quantification of soluble IL-22BP (E) and IL-22 (F) levels in the supernatant of gut biopsies from Ctrls ileum (n=4) and colon (n=12). (**G-H**) Comparison of IL-22 levels in the ileum (G) and the colon (H) of Ctrls and CD patients. (**I-J**) Comparison of IL-22BP levels in the ileum (I) and the colon (J) of Ctrls and CD patients. (**K-L**) Comparison of IL-22/IL-22BP ratios in the ileum (K) and the colon (L) of Ctrls and CD patients. Statistical significance for mean comparisons was assessed by the Mann-Whitney U-test in a two-sided manner, using a nominal significance threshold of P < 0.05.

Altogether, these results suggest that regional specialization in the intestine are characterized by higher levels of IL-22BP production in the ileum as compared to the colon in both controls and CD patients.

### The highest levels of IL22BP expression are detected in CD MNP and eosinophils

Various cell types can produce IL-22BP in IBD intestinal tissues, including populations of mononuclear phagocytes (MNP), which encompass dendritic cells (DCs) and macrophages, CD4^+^ T cells and eosinophils (14,15,19). So far, however, no systematic direct comparison of *IL22RA2* levels of expression by these different immune cell subsets has been performed specifically in intestinal tissues of CD patients. We thus analyzed *IL22RA2* expression in FACS-sorted populations of MNP (CD45^+^ HLA-DR^+^ CD11c^+^), eosinophils (CD45^+^ SIGLEC-8^+^), CD4^+^ T cells (CD45^+^ CD3^+^ CD4^+^), CD4^-^ T cells (CD45^+^ CD3^+^ CD4^-^), and B cells (CD45^+^ CD3^-^ CD19^+^) isolated from surgically resected intestinal tissues of CD patients (**Figures 2A&B, Supplementary Figure 1A**). Compared to the three lymphocytes fractions, MNP and eosinophils expressed higher levels of IL-22BP mRNA (**Figure 2B**). We observed similar trends between MNP and T cells isolated from CD mesenteric lymph nodes (MLN) (**Figures 2C&D, Supplementary Figure 1B**). Of note, similar mRNA levels were detected between populations of naïve and non-naïve T cells. Using IFI approaches, we previously verified IL-22BP protein expression in major basic protein (MBP)^+^ eosinophils and HLA-DR^+^ MNP in the gut LP but not in CD3^+^ T cells (14). We could not detect IL-22BP in CD3^+^ MLN cells either (**Supplementary Figures 1C**), hence possibly reflecting a lack of method sensitivity to capture lower expression levels in T cells. Finally, while we could observe increased expression levels of activation-induced *IL2RA* (encoding for CD25) in sorted gut and MLN T cells upon simulation with PMA and ionomycin, no difference existed for *IL22RA2* (**Supplementary Figures 1D-G**).

**Figure 2:**
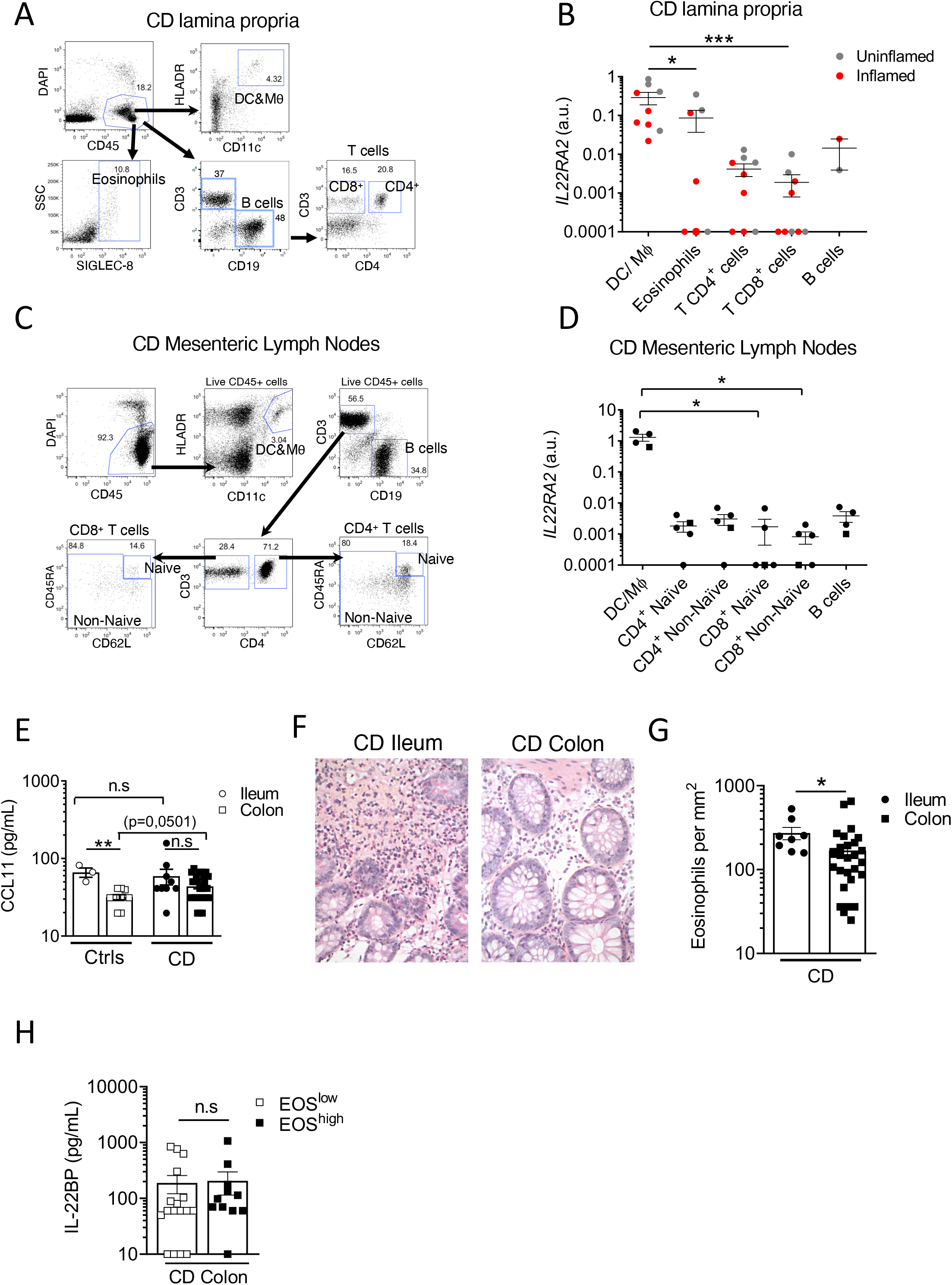
The highest levels of IL22BP expression are detected in CD MNP and eosinophils. **A**) Representative dot plots of the gating strategy used to sort indicated cell subsets from the lamina propria (LP) of intestinal surgical resections from Crohn’s disease (CD) patients. (**B**) *IL22RA2* expression was analyzed by RT-qPCR in indicated FACS-sorted cell subsets from CD patient gut LP samples (n=6). Grey and red dots refer to cells sorted from surgical resections respectively defined as non-inflamed and inflamed after pathology examination. (**C**) Representative dot plots of the gating strategy used to sort indicated cell subsets from mesenteric lymph nodes (MLN) of CD patients. (**D**) *IL22RA2* expression was analyzed by RT-qPCR in indicated FACS-sorted cell subsets from CD patient MLNs (n=5). (**E**) Levels of soluble CCL11 were quantified in gut endoscopic biopsy supernatants from controls (Ctrls) (n=12) and CD (n=37) patients after 6 hours of ex vivo culture. Dots and squares refer to ileum and colon tissues respectively. (**F**) Representative pictures of slides from formalin-fixed, paraffin-embedded (FFPE) sections of CD tissues stained with standard hematoxylin and eosin (HES) coloration. Original magnification x 200. (**G**) Eosinophil counts were scored in a blinded manner by a trained pathologist on HES slides from CD patients (n=36). Each count was obtained by averaging the results obtained from the analysis of different 5 fields and was expressed as eosinophils per mm^2^. (**H**) Comparison of soluble IL-22BP levels in the supernatant of endoscopic colonic biopsies of CD patients stratified into eosinophils (EOS)^high^ and EOS^low^ based on the median value. Mean comparisons of unpaired samples were performed using the Kruskal Wallis (for > 2 conditions) or the Mann-Whitney U-test. P-value < 0.05 were considered statistically significant.

Taken together, these data suggest that in a cell-intrinsic fashion, MNP and eosinophils express IL-22BP at the highest levels in intestinal tissues of CD patients.

### High levels of IL22BP in the ileum associate with higher proportions of eosinophils

In inflamed CD tissues, the strongest factor linked to IL-22BP variability was disease location (**Table 1**), and as for controls, IL-22BP secretion in biopsy supernatants was higher in the ileum than in the colon (**Figure 1**). Interestingly, the levels of CCL11 (aka. eotaxin-1), the main driver of eosinophil homeostatic recruitment in the intestine (22), were also higher in biopsy supernatants from the ileum than from the colon of controls (**Figure 2E**), concordant with the reported increased abundance of gut resident eosinophils in the small intestine (23– 25). This suggested that the higher levels of IL-22BP we detected in inflamed CD ileums as compared to CD colons could in part be explained by the heterogenous distribution of resident-eosinophils across intestinal segments. Accordingly, we confirmed the existence of higher eosinophil numbers in CD ileums than in CD colons (**Figures 2F&G**). Blood-derived eosinophil infiltration has been reported during colonic inflammation, including in CD (26–28), and CCL11 levels were indeed moderately increased in CD colons as compared to controls (**Figure 2E**). Median-based stratification of colonic CD patients into eosinophil^high^ and eosinophil^low^, however, did not reveal differences in IL-22BP production between the two subgroups (**Figure 2H**). Importantly, we previously reported that *IL22RA2* expression was undetectable in human peripheral blood eosinophils (14). This suggested that the contribution of eosinophil-derived IL-22BP detected in CD lesions could in fact reflect a pre-established production by tissue-imprinted gut-resident eosinophils but not by recently recruited eosinophils in inflamed tissues.

In consequence, our interpretation of these data is that homeostatic enrichment of tissue-resident eosinophil contributes to create an IL-22BP-rich environment that is maintained upon inflammation induction in CD ileums and participate to limit the extent of IL-22 bioavailability.

### IL22BP levels correlate with CCL24 produced by monocyte-derived dendric cells in the colon of CD patients

In the colon of CD patients, IL-22BP detected in biopsy supernatants spanned a wide range of concentrations, and while means did not differ from controls, a subset of patients nevertheless exhibited levels reaching those observed in the ileum (**Figures 1B**). As described above, this variability could not be explained by colonic eosinophil abundance (**Figure 2H**), which suggested a role for other IL-22BP sources (i.e. MNP and/or T cells). We were unable to examine IL-22BP production by flow cytometry because in our hands none of the anti-IL-22BP antibodies commercially available could be confidently validated. To infer possible sources, we hypothesized that IL-22BP secretion should be correlated with other cytokines in biopsy supernatants, in part as a consequence of shared cellular origins. We characterized the cytokine milieu associated with IL-22BP secretion in the CD colon through a multiplex analysis on the same biopsy supernatants as used for IL-22BP quantification. We then realized a correlation matrix to identify possible cytokines sharing similar regulation of expression as IL-22BP (**Figure 3A**). The strongest correlate of IL-22BP secreted levels was CCL24 (aka. eotaxin-2; spearman r=0,7; P<0,0001). Accordingly, the median-based stratification of CD patients based on their CCL24 levels in the colon showed that almost all CCL24^low^ patients had IL-22BP levels below the lower limit of quantification, while it was the opposite for CCL24^high^ patients (**Figure 3B**). We thus compared *CCL24* expression between T cells and MNP isolated from the CD lamina propria. The highest levels were detected in MNP (**Figure 3C**), which suggested they could be the privileged source responsible for IL-22BP heterogeneous secreted levels across colonic CD patients.

**Figure 3:**
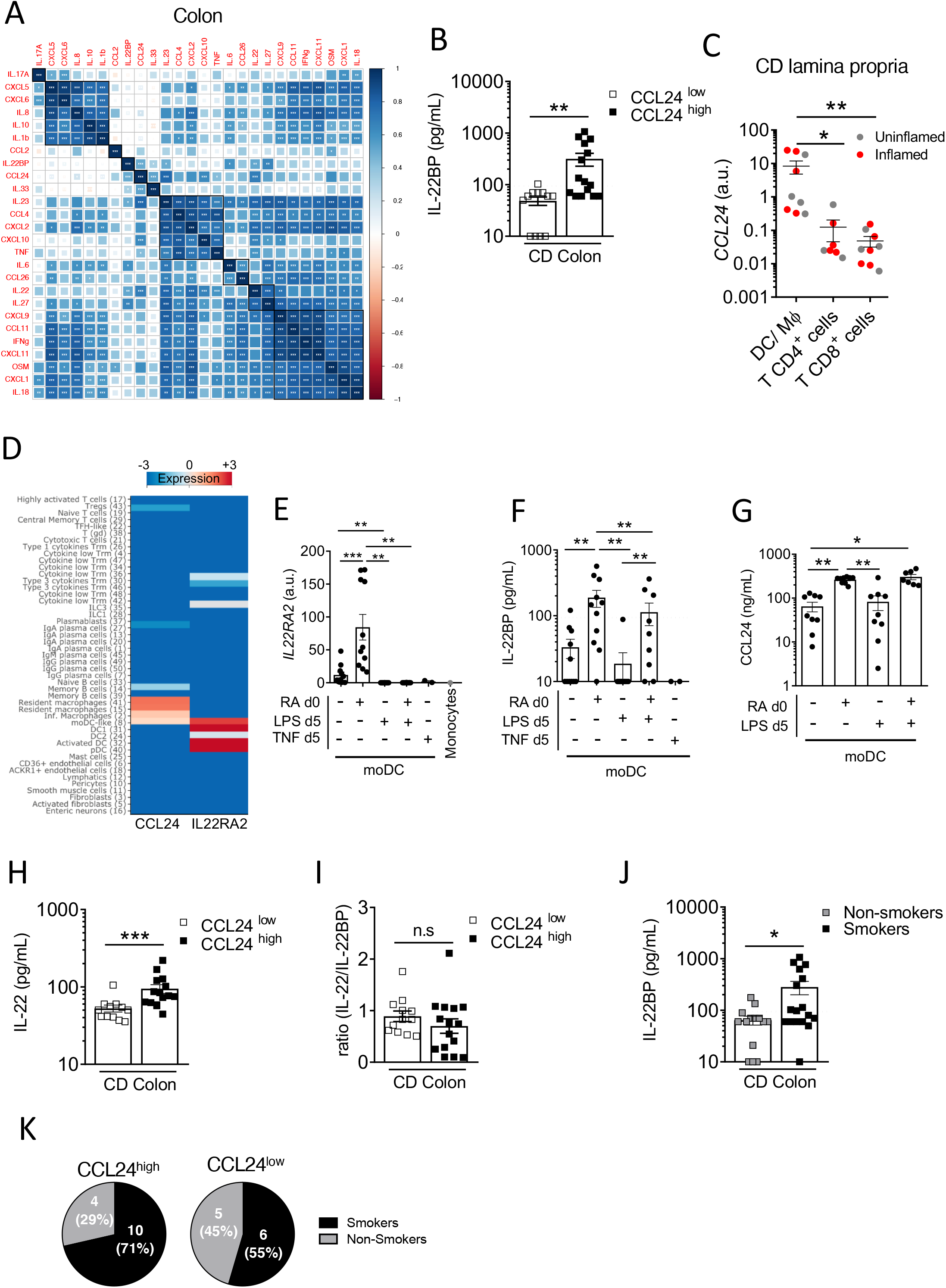
IL22BP levels correlate with CCL24 produced by monocyte-derived dendric cells in the CD colon. (**A**) Spearman correlation and hierarchical clustering of indicated cytokines quantified in culture supernatants of endoscopic colonic biopsies from 28 CD patients. (**B**) Comparison of soluble IL-22BP levels in CD patients stratified into CCL24^low^ and CCL24^high^ patients based on the median level. (**C**) *CCL24* expression was analyzed by RT-qPCR on FACS-sorted cells from CD lamina propria, as in Figure 2A. (**D**) Heatmap representing the expression of *IL22RA2* and *CCL24* in indicated cell types using the clustering analysis of a previously published single-cell RNA-sequencing (scRNA-seq) dataset (Martin JC, Cell, 2019). (**E-G**) Human blood classical monocytes from healthy donors were differentiated into monocyte-derived DC (moDC) with GM-CSF and IL-4 for 6 days in the presence of indicated ligands. At day 6 of culture *IL22RA2* mRNA expression was analyzed by RT–qPCR in moDC (E), and levels of IL-22BP (F) and CCL24 (G) were quantified in culture supernatants. (**H-I**) Comparison of soluble IL-22 (H) and IL-22/IL-22BP ratios (I) in culture supernatants of colonic CD biopsies from CCL24^high^ or CCL24^low^ patients. (**J**) Comparison of soluble IL-22BP levels in culture supernatants of colonic CD biopsies (n=32) from patients stratified according to their smoking status. (**K**) Pie charts representing the proportion of active smokers vs. non-smokers in the colon of CCL24^high^ or CCL24^low^ CD patients.

In an attempt to further refine the nature of a possible MNP subset producing both IL-22BP and CCL24, we reanalyzed a CD single-cell RNA-sequencing (scRNA-seq) dataset we published recently (29). While the contribution of eosinophils could not be evaluated in these data because of scRNAseq technical limitations, our analysis confirmed the highest expression levels of *IL22RA2* were present in MNP populations of DCs followed by lymphocyte populations including Type 3 cytokine-producing T cells, while *CCL24* was preferentially detected in macrophages (**Figure 3D**) (see (29) for detailed description about cluster annotations). Interestingly, only monocyte-derived dendritic cells (moDC) displayed shared detectable expression of both genes (**Figure 3D**), and we validated that moDC differentiated *ex vivo* co-secreted IL-22BP and CCL24 (**Figures 3E-G**). Importantly, we previously showed that retinoic acid (RA) was a potent inducer of *IL22RA2* expression in moDC (16), which we could further confirm at the protein level in moDC supernatants (**Figures 3E&F**). Rather interestingly, the same held true for CCL24 (**Figure 3G**), suggesting IL-22BP and CCL24 inductions could share similar regulatory pathways, such as RA activity in moDC. In turn, the activation of moDC with LPS for 24h had no effect on CCL24 secreted levels but decreased the production of IL-22BP (**Figures 3E&F**), hence supporting ours and others previous observations (16,30).

Other myeloid-derived cytokines showed significant positive correlations with IL-22BP, though weaker than for CCL24 (Spearman; P <0,05) (**Figure 3A**). These included IL-6, IL-27 and IL-23, the most potent inducer of IL-22 in CD4^+^ T cells and ILC3 (31,32). Concordantly, IL-22 correlated with IL-22BP and was also increased in CCL24^high^ patients, hence leading to similar IL-22/IL-22BP ratios between the two subgroups of CCL24^high^ and CCL24^low^ patients (**Figures 3H&I**). The co-regulation of IL-22/IL-22BP secretion in the colon of CD patients was in agreement with previous results we and others reported in IBD (14,15). Because CD4^+^ T cells also produce IL-22BP in IBD tissues (15), it is thus possible that Type 3 cytokine-producing T cells contribute to increase both IL-22 and IL-22BP levels in the colon of CD patients.

### Higher levels of IL-22BP are detected in the colon of actively smoking CD patients

Our multiple linear regression analysis suggested tobacco smoking as a possible additional factor associated with IL-22BP production in CD, though not reaching statistical significance (Student t-test t=1.813; P=0.08) (**Table 1**). We thus analyzed IL-22BP secretion in the colon of CD patients based on their smoking status. The production of IL-22BP was significantly higher in actively smoking patients (**Figure 3J**). A role for tobacco smoking has been suggested for the reprogramming of macrophages into the production of more immunoregulatory molecules, including CCL24 among others (corresponding to *in vitro* so-called “M2” macrophages) (33–35). Because a strong association existed between IL-22BP and CCL24 secretions in CD colons, we compared the proportions of smokers and non-smokers between CCL24^high^ and CCL24^low^ CD patients and verified an enrichment of smokers in the CCL24^high^ subgroup, though not reaching statistical significance (**Figure 3K**). While this would deserve further investigations, it is thus possible that tobacco-derived metabolites be part of the signals shaping moDC toward higher production of IL-22BP and CCL24 in the CD colon.

## Discussion

Our understanding of the pathophysiological functions assumed by the IL-22/IL-22BP axis in IBD is currently described in a rather homogenous way, frequently intermixing conclusions obtained from mouse and human studies, and including results generated in both CD and UC. By providing new insights focused on a more detailed characterization of IL-22BP biology in tissues from CD patients specifically, our study unravels a more complex picture arguing against a such generalization. Our work reveals that the distribution of IL-22BP production is not homogenous between CD patients and we suggest this could be the product of multiple factors related to IL-22BP complex biology in the human gut. First, we show that IL-22BP levels are higher in inflamed ileums when comparing to inflamed colons of CD patients, and we suggest this could be explained by distinct homeostatic distributions of gut resident-eosinophils between the two segments. This is of particular relevance as the injection of recombinant IL-22 is currently under evaluation in IBD clinical trials (NCT03650413), and it is thus unclear whether the IL-22BP-rich environment created in the ileum could lead to more interference with the biological actions of the drug in involved ileum vs. colon of CD patients. In addition, our data suggest that a raise of IL-22 bioactivity is more likely to be achieved in inflammatory responses developing in the colon, hence indicating that greater clinical benefits of IL-22 injections may nevertheless be expected in involved ileums of CD patients. More human studies will be needed to address this important question, as we previously showed that lL-22BP expression in gut resident-eosinophils is a feature not conserved in rodents (14). The reason why IL-22 is not induced in inflamed ileums despite higher homeostatic production in control tissues would also deserve more investigations. Beyond the difference of IL-22BP production created by regional specializations across gut segments, we also uncover additional heterogeneity in inflamed colons of CD patients. In particular, we observe a strong correlation between IL-22BP and CCL24 secretion and suggest this could be the consequence of a shared production by moDC with variable intensity across subsets of patients. Importantly, while, CCL24 was initially described as an eosinophil chemoattractant trough the receptor CCR3, experiments in rodents suggested it was not involved in eosinophil infiltration during colitis (28,36). Our data further suggest that IL-22BP variability among inflamed CD colons was likely not explained by the level of infiltration of blood-derived eosinophils but relied more on other IL-22BP cellular sources such as moDC. CCR3 is also expressed by subsets of T cells (37) and it is possible that CCL24-mediated T cell infiltration participates to the total production of IL-22BP in the colon of CD patients (15). Supporting the existence of similar regulatory pathways for both IL-22BP and CCL24 in inflamed CD colons, we report that retinoic acid increases the secretion of both proteins by moDC. As suggested by the higher IL-22BP levels we noticed in tobacco smokers, it is likely that additional factors contribute to differential regulations of IL-22BP production across patients. In that regard, it is interesting to note that IL-22BP levels also correlated with those of IL-22 and IL-23, the main driver of IL-22 expression in lymphocytes. Recent evidence suggests that lymphotoxin (LT)α1β2 upregulates IL22BP via the noncanonical NF-κB pathway in moDC (19). Because NF-κB is involved in the generation of IL-22-producing T cells, in part indirectly through the induction of IL-23 secretion in MNP (38), one may speculate that privileged cellular circuits exist in inflamed colons of a subset of CD patients, in which LTα1β2 from CD4 T cells drives both IL-22BP and IL-23 expression in moDC (39,40). The latter in turn can promote IL-22 production in CD4 T cells. While this would deserve further investigations, specific cellular niches in subsets of CD patients could thus participate to increase both IL-22 and IL-22BP global levels associated with a more localized impact on epithelial cell biology (19, 41). Clarifying the existence, tissue distribution and pathophysiological actions of such cellular circuits would be especially relevant in light of the upcoming IL-23 blockers in the CD therapeutic armamentarium (42,43).

In conclusion, our work provides new important information about the biology of IL-22BP in CD, which can open the way to therapeutically relevant future studies with regard to the modulation of the IL-22/IL-22BP axis in patients.

## Acknowledgements

This authors warmly thank Dr Mathieu Uzzan (Hopital Henri Mondor, Assistance Publique-Hôpitaux de Paris) for his critical reading of the manuscript and meaningful comments. This work was supported by grants Association Francois Aupetit (AFA), the European Crohn and Colitis Organization (ECCO), and Région Pays de la Loire (INSET) to RJ. JCM is supported by NExT “Junior Talent” and ANR JCJC (ANR-20-CE17-0009). AF was supported by a fellowship from CHU de Nantes (“Année supplémentaire d’Internat”).

## Author contributions

JCM, RJ and AF designed experiments and interpreted the data; JCM wrote the manuscript; AF, RJ and JCM edited the manuscript; AF, EL, TL, LD, SB and GB performed experiments and analyses; CB and NS performed cytokine multiplex assays; EM and AM performed *in vitro* moDC experiments; JP and AB managed the biobank and selected the patients included in the study.

## Data availability statement

The original contributions presented in the study are included in the article/supplementary material, further inquiries can be directed to the corresponding authors at the following e-mail addresses

---

**Supplementary Figure 1.**
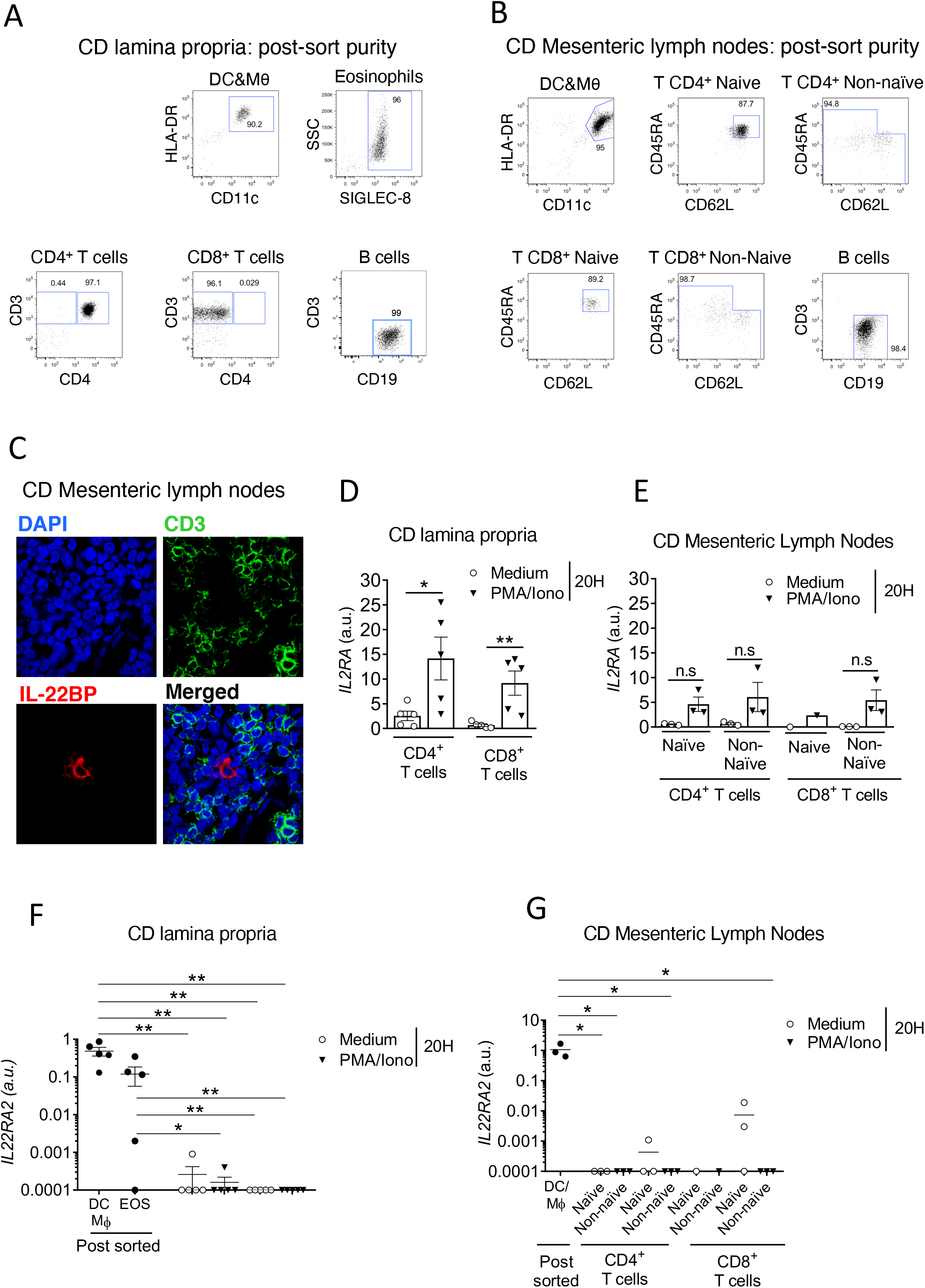
(**A-B**) Representative dot plots showing the post-sort purity of indicated cell subsets sorted from the lamina propria of intestinal surgical resections (A) and mesenteric lymph nodes (MLN) (B) of CD patients. (**C**) Representative IFI pictures of slides from formalin-fixed, paraffin-embedded (FFPE) sections of CD MLN stained with mAbs against IL-22BP (red) and CD3 (green), as well as with DAPI (blue) (n=3 CD patients). Original magnification x 630. (**D-G**) Indicated T cell subsets were sorted from CD lamina propria intestinal surgical resections (n=5) (D, F) and MLN (n=3) (E, G) and cultured with or without PMA (50ng/mL) and Ionomycin (250ng/mL) (PMA/Iono). After 20 hours, expressions of *IL2RA* (D-E) and *IL22RA2* (F-G) were analyzed by RT-qPCR. IL22RA2 expression levels of freshly isolated DC/Macrophages and eosinophils are indicated for reference. Means comparisons were performed with the Kruskal– Wallis test for unpaired samples and with the Wilcoxon test for paired samples. P-value < 0.05 were considered statistically significant. CD: Crohn’s disease; DC: dendritic cells, EOS: eosinophils; Mθ: macrophages.

