## Supplemental Table 1 for "IL-22BP production is heterogeneously distributed in Crohn’s disease"

**Supplementary Table 1: Clinical characteristic of controls and Crohn's disease patients cohort 1**

| **Characteristic** | **Controls** | | **Crohn's disease Cohort 1** | |
|---|---|---|---|---|
| **Samples** | *Ileum* | *Colon* | *Ileum* | *Colon* |
| **General data** | n=4 | n=12 | n=11 | n=32 |
| Sex (F/M) | 3/1 | 7/5 | (8/3) | 17/15 |
| Age (years)* | 54 ± 11.7 | 64 ± 11.9 | 35 ± 6,7 | 35 ± 12,4 |
| Disease evolution (years)* | - | - | 13 ± 9,3 | 9,4 ± 7,5 |
| Smokers (yes/no) | - | - | (2/9) | (17/15) |
| **Montreal classification** | | | | |
| **Age at diagnosis** | | | | |
| *A1 below 16 y.o* | | | 1 | 3 |
| *A2 between 17 and 40 y.o* | | | 10 | 29 |
| *A3 above 40 y.o* | | | 0 | 0 |
| **Location** | | | | |
| *L1 ileal* | | | 6 | 4 |
| *L2 colonic* | | | 0 | 4 |
| *L3 ileocolonic* | | | 5 | 24 |
| *L4 isolated upper disease* | | | 0 | 0 |
| **Behavior** | | | | |
| *B1 non-stricturing/penetrating* | | | 6 | 9 |
| *B2 stricturing* | | | 2 | 10 |
| *B3 penetrating* | | | 2 | 6 |
| *B2&B3* | | | 1 | 7 |
| **Medications at time of endoscopy** | | | | |
| *5-Aminosalicylic acid (5-ASA)* | | | 0 | 1 |
| *Anti-TNF* | | | 5 | 19 |
| *Immunosuppressants (IS)* | | | 4 | 7 |
| *Corticosteroids* | | | 0 | 0 |
| *Anti-integrin* | | | 0 | 5 |
| *None* | | | 4 | 3 |

*mean±sd,
