## Supplemental table 2 for "IL-22BP production is heterogeneously distributed in Crohn’s disease"

**Supplementary Table 2: Clinical characteristic of Crohn's disease patients cohort 2**

| Characteristic | Crohn's disease Cohort 2 |
|---|---|
| **General data** | n=6 patients |
| Sex (F/M) | (4/2) |
| Age (years)* | 51.7 ± 20.6 |
| Disease evolution (years)* | 14.4 ± 16.3 |
| Smokers (yes/no) | (n.d for 1; 0/5) |
| **Montreal classification** | |
| **Age at diagnosis** | |
| *A1 below 16 y.o* | 1 |
| *A2 between 17 and 40 y.o* | 2 |
| *A3 above 40 y.o* | 3 |
| **Location** | |
| *L1 ileal* | 1 |
| *L2 colonic* | 2 |
| *L3 ileocolonic* | 3 |
| *L4 isolated upper disease* | 0 |
| **Behavior** | |
| *B1 non-stricturing, non-penetrating* | 0 |
| *B2 stricturing* | 3 |
| *B3 penetrating* | 2 |
| *B2&B3* | 1 |
| **Medications at time of surgery** | |
| *5-Aminosalicylic acid (5-ASA)* | 1 |
| *Anti-TNF* | 2 |
| *Immunosuppressants (IS)* | 2 |
| *Anti-IL-23* | 1 |
| *Corticosteroids* | 2 |
| *None* | 0 |

*mean±sd, n.d : not done
